# Proteostasis Associated Mutations in HSPB5 Destabilize the αB-crystallin Domain

**DOI:** 10.1101/2024.11.12.623266

**Authors:** Mst. Shanzeda Khatun, Md. Chayan Ali, Largess Barua, Yeasmin Akter Munni, Raju Dash, Md Mafizur Rahman

**Author notes:** **Corresponding Author: Md Mafizur Rahman,** Department of Biotechnology and Genetic Engineering, Faculty of Biological Sciences, Islamic University, Kushtia-7003, Bangladesh; **Raju Dash,** Department of New Biology, Daegu Gyeongbuk Institute of Science & Technology (DGIST), DGIST, E4-620, 333, Technojungang-daero, Hyeonpung-myeon, Daegu, 42988, Republic of Korea. Equal contribution (Mst. Shanzeda Khatun, Md. Chayan Ali).

## Abstract

HSPB5 (αB-crystallin) is an ATP-independent, stress-inducible chaperone protein that improves protein misfolding and degradation in proteotoxicity-related diseases, including several myopathies, neurodegenerative diseases, and cancers. Although single nucleotide polymorphism (SNP) reduces the overall activity of HSPB5, its dynamic behaviour remains unknown. To get molecular insights into the deleterious, pathogenic, and proteotoxicity-related mutations, this study investigates the potential deleterious SNPs associated with HSPB5. Notably, eleven computational tools identified D109H, R120G, and D140N as the most deleterious SNPs from a total of 313 missense SNPs. Interestingly, these three mutations are present in the core αB-crystallin (αB-c) domain. A molecular dynamics simulation for 500 ns was conducted to reveal these variants’ mechanistic insights. The mutant variants showed higher flexibility and significant conformational changes than the wild, which might be noteworthy to reduce these variants’ chaperoning activity. Also, this conformational change elucidated the loss of function mutations, which could alter these variants’ oligomeric properties. This study will help our understanding of the role and molecular mechanism of HSPB5 mutations in proteotoxicity vulnerable diseases.

## 1 Introduction

Proteostasis (protein homeostasis) is the balance between protein synthesis and folding to degradation. The molecular and cellular processes are dependent on proteostasis. Proteostasis failure is associated with various diseases, including neurodegeneration [1]. The proteostasis network (PN) is a complex signaling cascade that regulates the cellular location, amount of protein synthesis, and assembly of different polypeptides at a time. Besides, PN also prevents protein misfolding and accumulation [2]. The molecular chaperones are the crucial factors of PN, which regulate the correct folding, conformational stability, and clearance of misfolded proteins [2, 3]. In this network, the ubiquitous small heat shock (sHSP), alpha-basic-crystallin (αB-c) plays a significant role, including’s regulation of caspase activity, prevent denatured protein accumulation, actin, and titin polymerization, control cellular redox state, maintain cytoskeleton integrity, and proteasomal degradation [4]. The αB-c, also called HSPB5, a ∼20 kDa protein produced from the *CRYAB* gene and found highly in eye lens (maintains lens transparency) and heart muscle and acts as a molecular chaperone. The human HSPB5 contains three domains, the core αB-c domain (ACD), which is fringed by N and C terminal domains [5, 6], and found to express in skin, muscle, kidney, lung, spleen, and brain and associated with cardiac myopathies and neurodegenerative diseases (NDD) [7, 8]. The large oligomeric forms of sHSPs exchange their subunit, which is supposed to act in chaperoning function [9]. HSPB5 acts on various effectors and regulates apoptosis. It was reported that the upregulation of HSPB5 in rabbit lens epithelial cells prevents activation of RAS and inhibits RAF/MEK/ERK signaling, which eventually decreases apoptosis [10]. Besides, HSPB5 protects cells from various stresses like oxidative stress, inflammatory stress and inhibits caspase-3, PARP, and nuclear translocation of Bax and Bcl-2 [11]. Overexpression of HSPB5 also plays a neuroprotective role in NDD, such as familial amyloidotic polyneuropathy [11]. It is also found that the core αB-c domain inhibits protein accumulation and amyloid toxicity [12]. Interestingly, individual ACD interacts with the sHSP oligomer to decipher chaperoning functions [9].

However, the mutation in HSPB5 protein hampers proteostasis and found an association with several diseases. For example, mutations in the ACD domain (D109H, R120G, D140N) [13, 14] are linked with cataracts, myopathy, myofibrillar myopathy, and cardiomyopathy [15–17]. The D109H variant of human HSPB5 was found to increase protein accumulation in HeLa cells [18], decline protein stability, abolish inter-domain residual interaction, decrease substrate contact [19], and reduce chaperone activity [20]. The R120G mutation abolishes substrate binding capacity [21, 22] and diminishes chaperone activity [21]. The loss of chaperone activity was also shown in D140N mutation [22], which similarly causes protein aggregation [18] and reduces substrate specificity [22]. These mutations were also shown to decrease the chaperone’s thermal stability and oligomerization, which reduce the protein’s overall functions [19, 22]. To our knowledge, the detailed molecular insights, i.e., structural behaviour, protein dynamics, conformational dynamics of HSPB5 due to mutational perturbance, have not been explored yet.

Therefore, the *in silico* study was aimed to detect the most pathogenic SNPs and insights into their damaging effects, emphasizing chaperone-regulated diseases through molecular dynamics simulation of the wild and mutant variants. Specifically, we explored the structural consequences of the most influential protein aggregate causing variants D109H, R120G, and D140N of the HSPB5 ACD domain.

## 2 Methodology

### 2.1 Collection of CRYAB SNP dataset

The *CRYAB* (HSPB5) gene missense SNPs datasets were repossessed from the “National Center for Biotechnology Information (NCBI) dbSNP (https://www.ncbi.nlm.nih.gov/snp/)” [23] database. The total methodology is depicted in **Fig. 1**.

**Fig 1:**
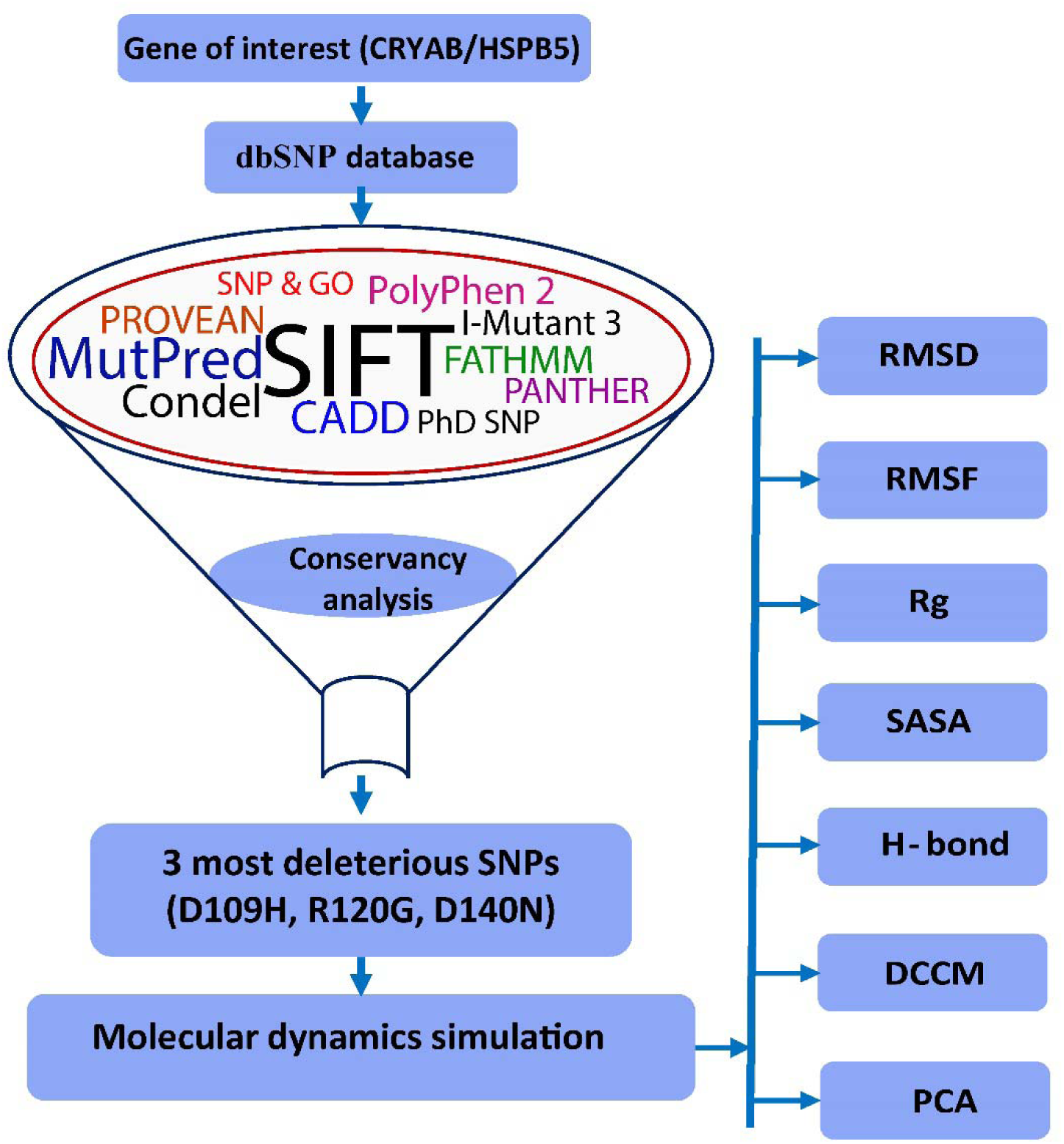
Schematic illustration of the followed schema of this study.

### 2.2 Identification of most pathogenic SNPs

To identify the most damaging *CRYAB* (HSPB5) SNPs, eleven most widely used tools were used i.e., Sorting Intolerant From Tolerant (SIFT) [24], Polymorphism Phenotyping v2 (PolyPhen-2) [25], Condel [26], Combined Annotation Dependent Depletion (CADD) [27], MutPred [28], Predictor of Human Deleterious Single Nucleotide Polymorphisms (PhD-SNP) [29], Protein Analysis Through Evolutionary Relationships (PANTHER) [30], Single Nucleotide Polymorphisms and Gene Ontology (SNP & GO) [31], Functional Analysis through Hidden Markov Models (FATHMM) [32], Protein Variation Effect Analyzer (PROVEAN) [33], and I Mutant 3.0 [34]. SIFT uses the sequence homology method and PSI-BLAST algorithm to predict deleterious effects [24], while PolyPhen v2 uses the Bayesian classifier and position-specific independent counts (PSIC) scoring techniques. In PolyPhen-2, two integrated algorithms were used; HumDiv and HumVar, in which HumDiv categorizes less damaging SNPs via PSIC, and HumVar classifies the extreme phenotypes [35]. Condel predicts deleterious SNP using an integrated approach of LogRE (Log R Pfam E value) [36]. CADD considers SIFT and PolyPhen derived protein level scores to predict deleterious mutations [27], whereas MutPred forecasts 14 different functional and structural properties of proteins either gain or losses due to mutation [28]. Both PhD-SNP and PANTHER use support vector machine (SVM) algorithms to identify deleterious or disease-causing amino acid (AA) substitutions. In addition, PhD-SNP uses the BLAST algorithm and UniRef90 database [29, 37]. SNP & GO classify disease-causing AA substitutions using the gene ontology database through molecular and functional properties [31]. FATHMM uses hidden Markov models (HMMs) to predict pathogenic mutations [32]. The sequence and alignment-based clustering scores are used by PROVEN to predict functional alteration due to mutation [33, 38]. Finally, I-mutant 3.0 uses SVM algorithms to analyze the stability of variants through Gibbs free energy changes [34].

### 2.3 Conservation analysis

The web-based conservation analysis tool ConSurf (http://consurf.tau.ac.il/) was used to analyze the evolutionary conservation of each amino acid position in a protein. The Bayesian calculation method was applied in this study [39, 40]. The conservation values of each quickly changing amino acid position vary; the amino acid positions that change slowly are conserved. The level of amino acid conservation is estimated based on a conservative 1-9 scale in which the most flexible positions are 1-3, the moderate positions are 4-6, and the highly conservative positions are 7-9 [41].

### 2.4 Simulation system preparation

The functional crystal structure of the ACD domain (PDB ID **2WJ7** [42]) of HSPB5 was extracted from the protein data bank (http://www.rcsb.org/pdb) [43]. The wild structure of HSPB5 was at first prepared through the addition of bond orders, hydrogen, and charges and polished by eliminating water atoms and adjusted at neutral pH. The structure was again adjusted by correction of some thiol groups, amine derivatives, hydroxyl groups, aspartic acids, protonated glutamic acids, and histidines. A minimization protocol was applied to fix heavy atoms Root Mean Square Deviation (RMSD) using OLPS3 (Optimized Liquid Simulation Potentials) force field until RMSD reached 0.30 Å. The substituted variants D109H, R120G, and D140N structures were generated using the Schrödinger Mutate Residues script 2017-1 suite (LLC, New York, NY, USA) through computational mutagenesis [44]. The structures of wild and D109H, R120G, and D140N variants were refined using short molecular dynamics (MD) simulations. The MD simulation was performed using YAMBER3 (Yet Another Model Building and Energy Refinement force field) [45] force field for 500 ps at 298 K, pH 7.4, and 0.997 g/cc solvent density.

### 2.5 Molecular dynamics simulation

The structural and dynamic behaviour of HSPB5 protein variants was analyzed by YASARA Dynamics (YASARA Biosciences GmBH, Austria) software [46] as described previously [47–49]. All selected structures were cleaned and optimized for hydrogen bonds. The AMBER14 force field was used in this simulation system [50]. Then, the system was solvated using the TIP3P (0.997 g/L density) water model. The titratable amino acids pKa values of all protein variants (acid dissociation constant) were calculated [51]. The TIP3P and AMBER14 force field provide the best outputs described as before [52–54]. Then, the system was neutralized by providing additional counter ions. A default salt concentration (0.9% Na+ Cl–) with 7.4 pH was maintained in the box. The simulated annealing protocol was used to diminish conformational stress using the steepest descent algorithm. The Particle Mesh Ewald method [55] was considered for long-range electrostatic interaction analysis and 8.0 Å cut off space interval for short-term electrostatic interactions, while uniform density approximation was used for long-range van der Waals (VDW) interactions. Finally, a 500 ns long simulation was carried out for each system with a Berendsen thermostat at a 2.0 fs time step interval along with multiple time-step algorithms [56]. At constant pressure, the trajectory was collected at every 25 ps. The YASARA suite integrated with preinstalled (md_run.mcr) macro was used in all simulation phases, and generated results were analyzed using VMD [57] and DSSP [58] tools. Lastly, the trajectories were used for RMSD; root mean square fluctuation (RMSF), radius of gyration (Rg), solvent accessible surface area (SASA), H-bond analysis, dynamic cross-correlation matrix (DCCM), and principal component analysis (PCA) analyses.

### 2.6 Dynamic Cross-Correlation Matrix

The inner conformational dynamics of proteins were shown by dynamic cross-correlation matrix analysis (DCCM). Here, the R program-oriented Bio3D [59] package was used. Bio3D DCCM offers correlation coefficients to the Pearson covariance matrices calling on “cov2dccm” and may calculate the following equation:

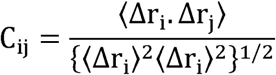

Where, Cα electrons are considered by the cross-correlation ratio, C_ij_, [60]. Δr_i_ and Δr_j_ signify the regular location of ith and jth residues, respectively, whereas the angular brackets show the time average. DCCM measured values fluctuated from −1 to +1, and positive and negative values denote positive and negative correlations, respectively.

### 2.7 Statistical Analysis

The correlation among all the tools was analyzed using the Microsoft Excel data analysis tools pack. The GraphPad Prism v8.0 (San Diego, USA) software carried out the MD trajectories’ statistical analysis considering the significant *p-*value, *p*<0.05.

## 3 Results

### 3.1 Selection, screening of dataset, and identification of deleterious SNPs

A total of 313 missense SNPs were initially considered for finding the most pathogenic mutations. The significant SNPs were identified with various computational tools represented in **Fig. 2B**. Among all mutations, missense variants were only 7.36 %, which was considered for further analysis (**Fig. 2A**). The cut off values of these tools were as follows SIFT (≤0.05), polyphen-2 HumDiv (LJ0.9), Polyphen-2 HumVar (LJ0.9), Condel (>0.9), CADD (>20), MutPred (>0.75), PhD SNP (>0.5), PANTHER (≥0.5), SNP & GO (>0.5), FATHMM (<-3.0), PROVEAN (<−2.5), and ILJMutant 3.0 (<−0.5). The highest deleterious SNPs were predicted by I-mutant 3 (203), shown in **Fig. 2B**. It is also shown that all the predicted SNPs in all algorithms were expressively correlated (*P*<0.0001) (**Fig. 2C**). The dark red color indicates a positive correlation. The result shows that D109H, R120G, and D140N variants are the most deleterious SNPs as predicted by all computational tools (**Table 1**). These mutations reduce the chaperone-like activity of HSPB5, and we anticipated the associated mechanisms involved.

**Fig. 2:**
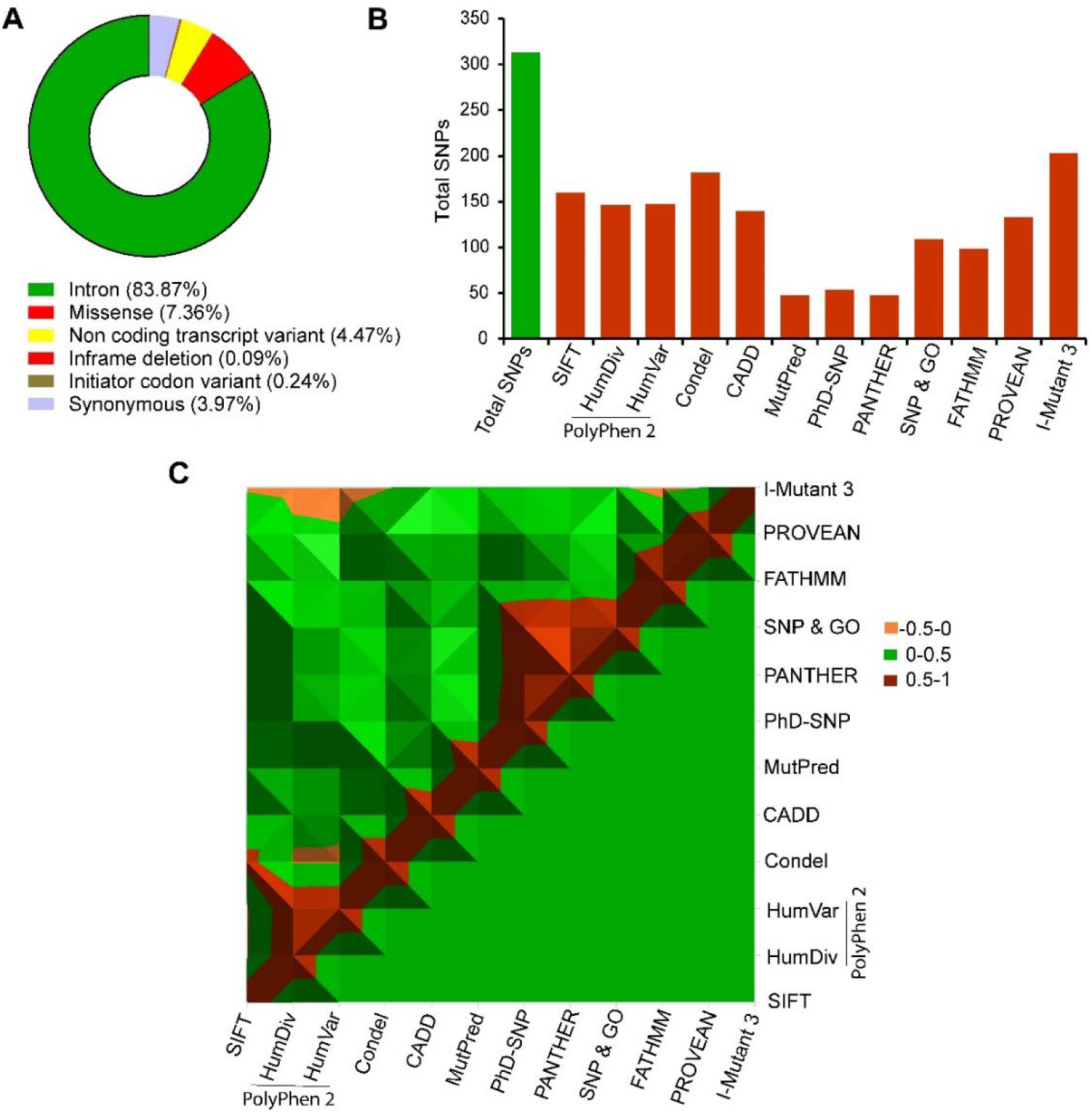
Distribution of SNPs where (A) Total variants class of *CRYAB* (HSPB5), (B) Total number of SNPs in the *CRYAB* gene is predicted by using various algorithms, and (C) The surface chart depicts the correlations amongst the predicted SNPs by all tools. The dark red color indicates a positive correlation.

**Table 1:** Most deleterious variants with their experimental evidence, disease association and in silico deleterious analysis.

| Variants | Experim<br>ental<br>evidence | Diseases<br>associati<br>on | Ref. | *In silico* SNP analysis |  |  |  |  |  |  |  |  |  |  |  |
| --- | --- | --- | --- | --- | --- | --- | --- | --- | --- | --- | --- | --- | --- | --- | --- |
|  |  |  |  | SIFT | PolyPhen 2 |  | Condel | CADD | MutPred | PHD | PANTH | SNP | FATHMM | PROVE | I Mutant-3 |
|  |  |  |  |  | HumDiv | HumVar |  |  |  | SNP | ER | &<br>GO |  | N |  |
| **D109H** | LCA,<br>PA, APT | CT,<br>CDM | [17, 61] | **0.00** | **1.00** | **0.998** | **0.911** | **34.00** | **0.766** | **0.832** | **0.892** | **0.932** | **-3.21** | **-6.16** | **-0.99** |
| **R120G** | LCA,<br>LSB,<br>CDM | CT,<br>CDM,<br>MA,<br>MM | [61] | **0.00** | **1.00** | **0.963** | **0.945** | **26.60** | **0.966** | **0.925** | **0.868** | **0.983** | **-3.68** | **-6.17** | **-2.18** |
| **D140N** | LCA, PA | CT,<br>CDM | [13, 61] | **0.00** | **0.999** | **0.994** | **0.892** | **28.30** | **0.803** | **0.642** | **0.668** | **0.897** | **-3.03** | **-3.95** | **-2.03** |

LCA.: Loss of chaperone activity; PA: Protein aggregation; APT: Apoptosis; LSB: Loss of substrate binding; CT: Cataract; CDM: Cardiomyopathy; MM: Myofibrillar myopathy; N/A: Not available. The scores in bold form indicate deleterious or damaging, according to the individual criteria.

### 3.2 Evolutionary conservancy analysis

The amino acids reside in the functional domain of proteins, or enzymes are conserved in nature and have the most vital role in substrate management to exert biological functions. Thus, nsSNPs, located in a highly conserved region of amino acid, evidently more harmful than nsSNPs located in non-converted positions [62]. Evolutionary information is essential for identifying mutations that could affect human health [63]. ConSurf uses an evolutionary variation to determine the conservation degrees in multiple sequence alignments. It defines functional and structural residues of a protein using two separate data, i.e., solvent accessibility and evolutionary conservation data. High conserved residues are either functional or structural depending on their location, whichever on the surface or inside the protein core [39]. The ConSurf result suggested that the D109H, R120G, and D140N residues are functional, highly conserved, and exposed (**Fig. 3**).

**Fig. 3:**
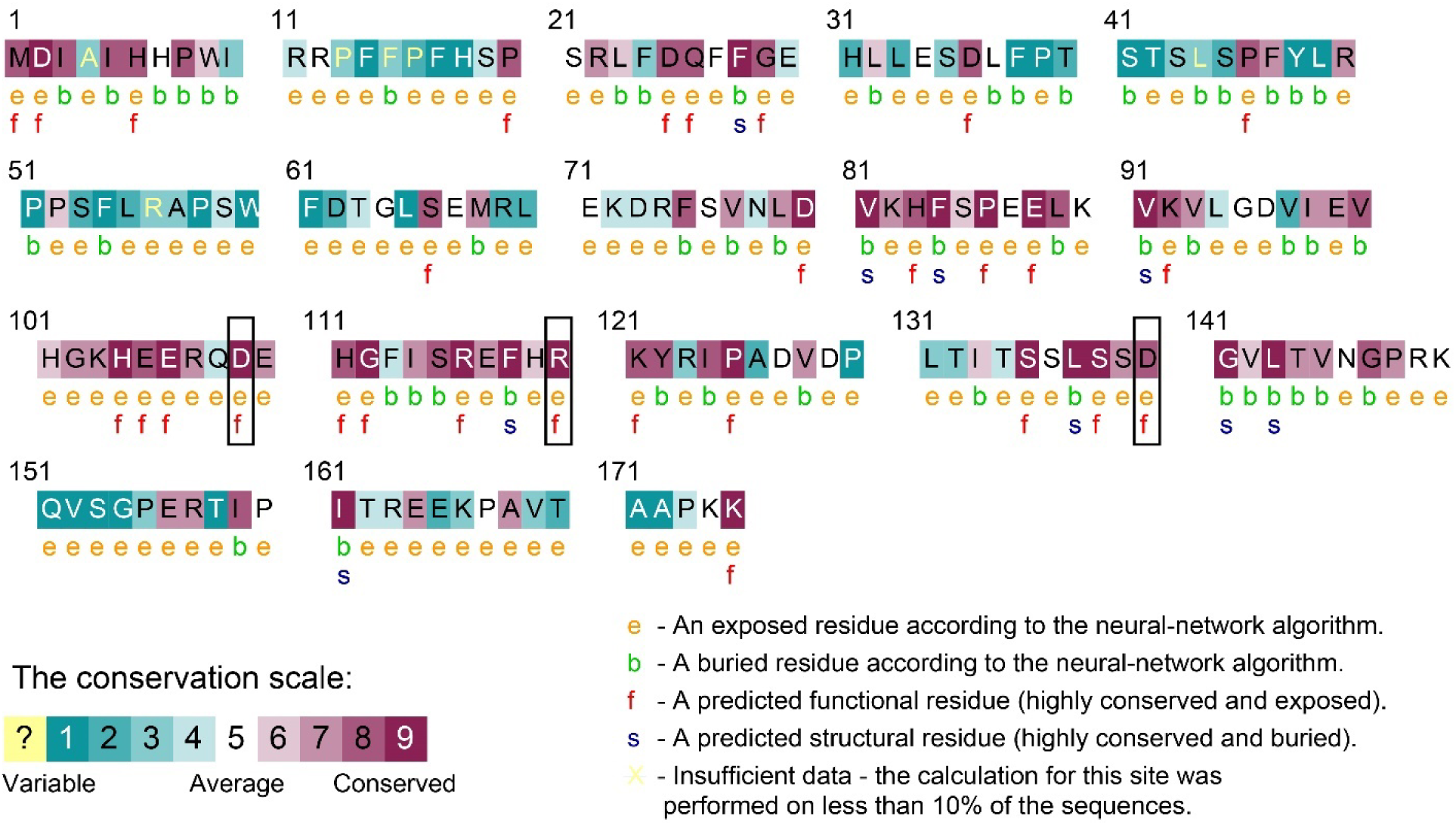
Conservation analysis. The black box indicates the most deleterious and highly conserved residues.

### 3.3 Molecular dynamics simulations

We employed the MD simulation method to understand the structural and functional behaviors of wild and mutant proteins at the atomic level. We applied different dynamics properties like RMSD, Rg, SASA, RMSF, H-bond, DCCM, and PCA analyses. In MD simulation, the RMSD demonstrates the conformational stability of proteins considering their backbone Cα atoms, as shown in **Fig. 4**. According to RMSD analysis, the wild variant achieved equilibrium at 25 ns and remained stable until 325 ns. The D109H variant achieved stable condition at 163 ns and remained stable until 463 ns (**Fig. 4A**).

**Fig. 4:**
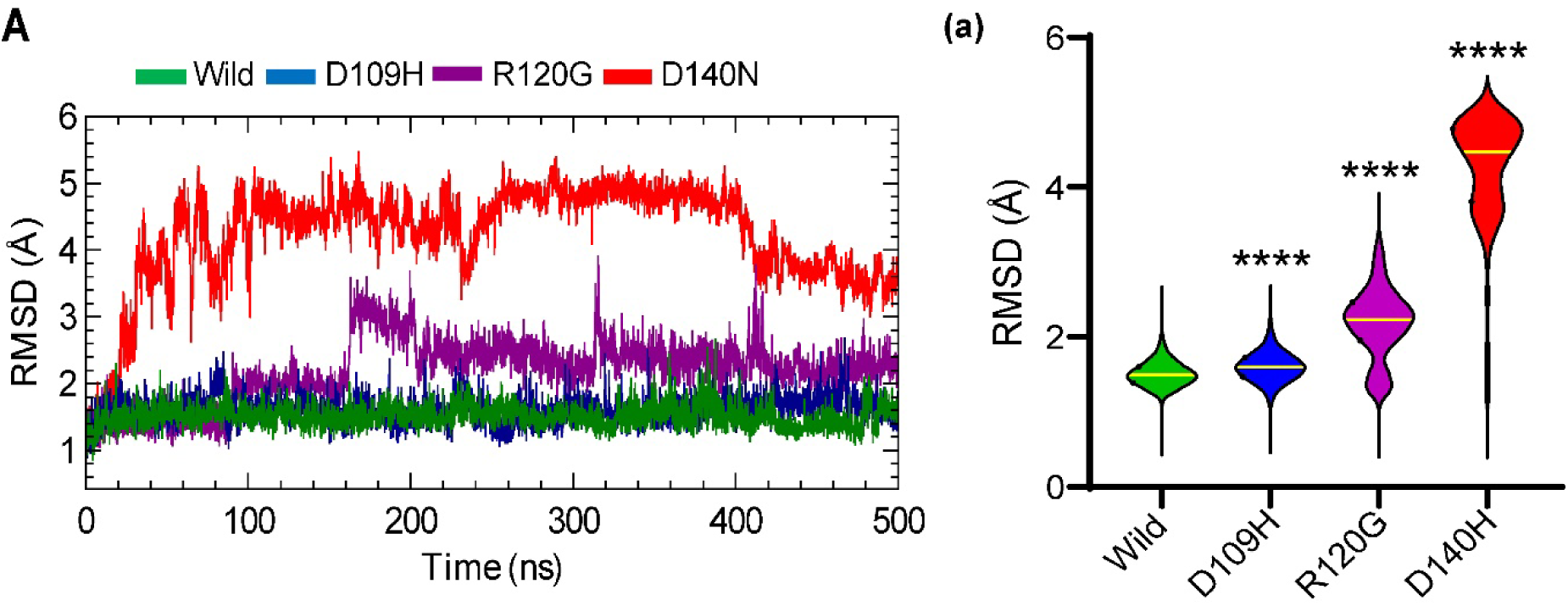
Conformational stability analysis. (A) Root mean square deviation analysis of all systems. (a) The violin plots indicate each system’s RMSD distribution and the statistical significance considering \*\*\*\**p*<0.05.

On the other hand, the R120G variant remained stable from 200 to 500 ns. The D140N variant showed an equilibrium state at 110 to 410 ns, and the RMSD decreased afterward, which indicates a significant conformational change of structures. Therefore, the RMSD results confer that mutation causes significant changes; thus, it warrants further investigation (**Fig. 4Aa**). For further study, the mentioned stable time frame of RMSD was considered.

#### 3.3.2 Effects in conformational dynamics

The radius of gyration (Rg) calculation was conducted to explore more detailed dynamic and conformational changes of mutant variants compared to the wild ones (**Fig. 5A**). Rg represents the global compactness of proteins considering the overall protein dynamics (higher Rg values indicate lower compactness) [64, 65]. **Fig. 5A** represents that mutation reduced the compactness of proteins. The averages Rg values are 14.82 Å, 15.20 Å, 15.01 Å, and 15.85 Å for wild, D109H, R120G, and D140N, respectively (**Table 2**). Overall, the Rg values denote that the identified mutation induces the flexibility of ACD domains compared to wild types (**Fig. 5Aa**).

**Fig. 5:**
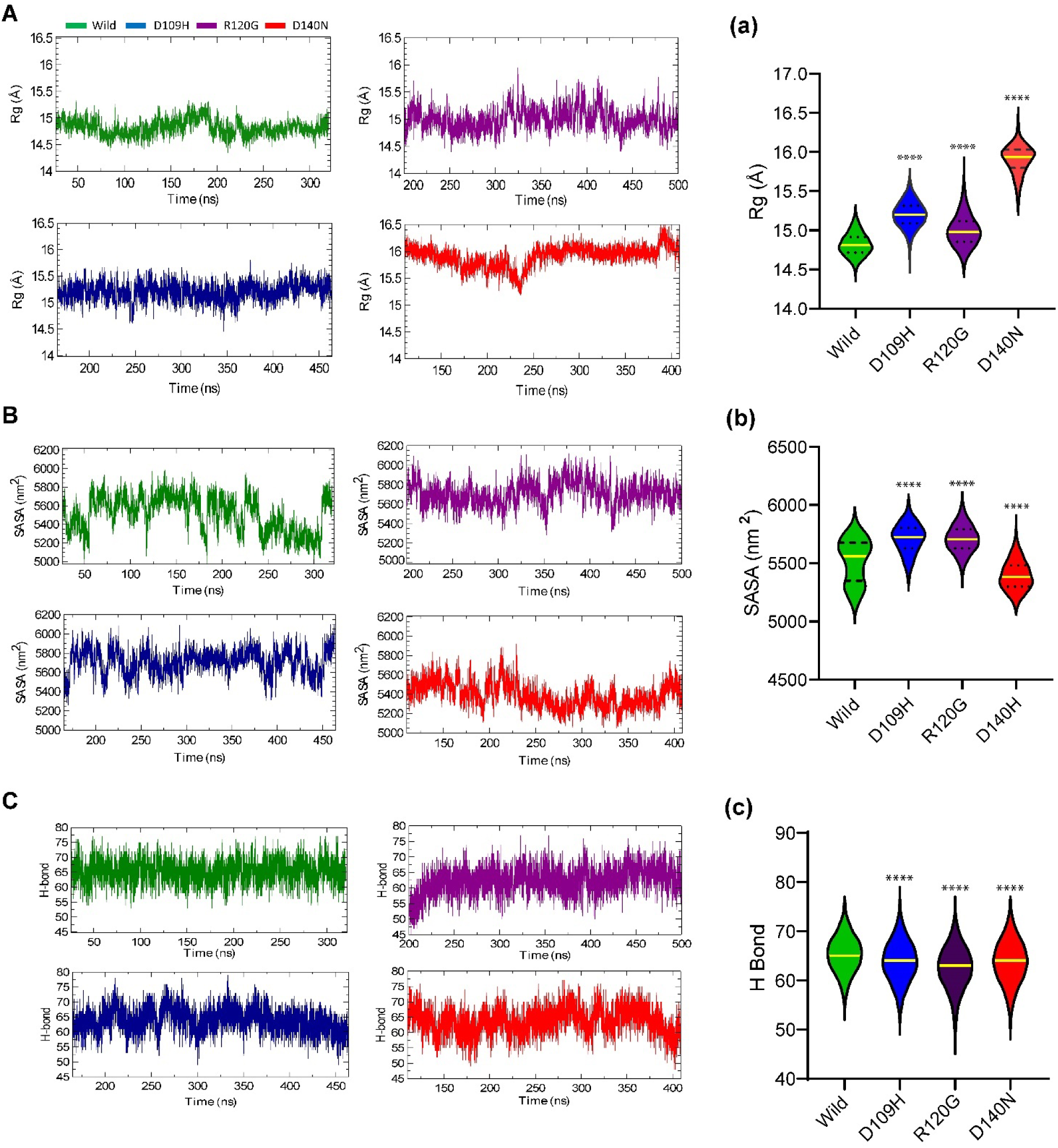
Effects of mutation on protein conformational stability of αB-crystallin domain in which (A) analysis of radius of gyration (Rg), (a) the violin plots indicate the Rg distribution of each system and also the statistical significance considering \*\*\*\**p*<0.05, (**B**) analysis of solvent accessible surface area (SASA), (b) The violin plots indicate the SASA distribution of each system and also the statistical significance considering \*\*\*\**p*<0.05. (C) analysis of total intra-residue hydrogen bond, and (c) the violin plot indicates the individual system’s distribution of total hydrogen bonds. In all cases, green, dark blue, purple, and red color represents wild, D109H, R120G, and D140N, respectively.

**Table 2:**
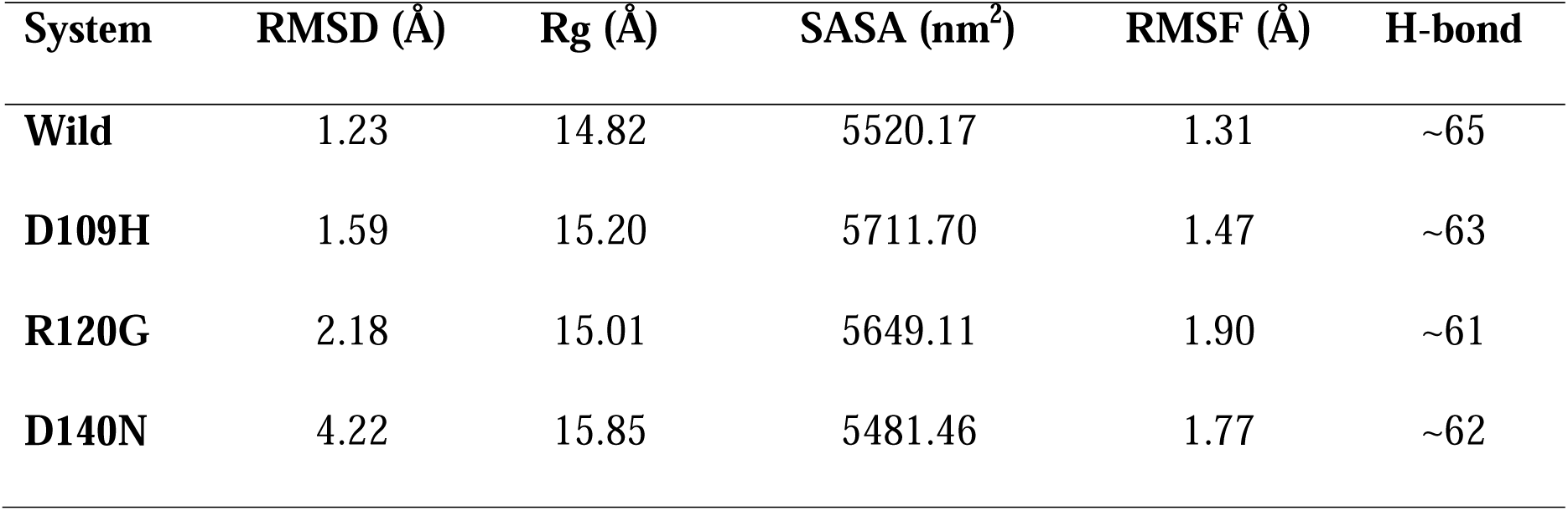
The average mean value of molecular dynamics simulation trajectories.

The solvent accessibility of these variants was measured by calculating SASA. The SASA values depict solvent accessibility, where lower SASA value denotes proteins’ contracted nature [66]. The SASA values of all variants are shown in **Fig. 5B**. The average SASA values of D109H (5711.70 nm^2^) and R120G (5649.11 nm^2^) are higher than the wild protein (5520.17 nm^2^), whereas D140N (5481.46 nm^2^) showed a lower average SASA value. The higher SASA values of D109H and R120G indicate their openness, whereas the reduced SASA value of D140N depicts decreased solvent accessibility (**Fig. 5Bb**). Overall, the changed solvent availability representing a modulated protein folding due to mutation.

The molecular interactions, i.e., hydrogen bonds, hydrophobic, and electrostatic interactions, determined proteins’ stability [67]. The protein stability is directly proportional to the number of interactions. The number of hydrogen bonds of wild and mutant variants is depicted in **Fig. 5C**. As shown in **Fig. 5C**, the number of average hydrogen bonds of wild protein is relatively higher than the mutant variants. The lower number of hydrogen bonds indicates the lower stability of these variants (**Fig. 5Cc**).

#### 3.3.3 Effects on protein dynamics

The local protein structure changes due to mutation were analyzed by the root mean square fluctuations (**Fig. 6**). The higher RMSF value denotes higher flexibility [66]. The violin plot (**Fig. 6A**) represents the statistical significance of RMSF values of the simulation data. The studied three mutants showed higher fluctuations throughout the whole simulations than that of the wild protein (**Fig. 6B**). The ACD domain of α -c contains two loops (loop 5/6) and six β strands. The D109H variant showed higher flexibility in the AA_92-95_ and AA_107-111_ residue regions (**Fig. 6B**). The D140N residue showed higher fluctuations in AA_106-112_, AA_115-124_, and AA_135-143_ residues (**Fig. 6B**).

**Fig. 6:**
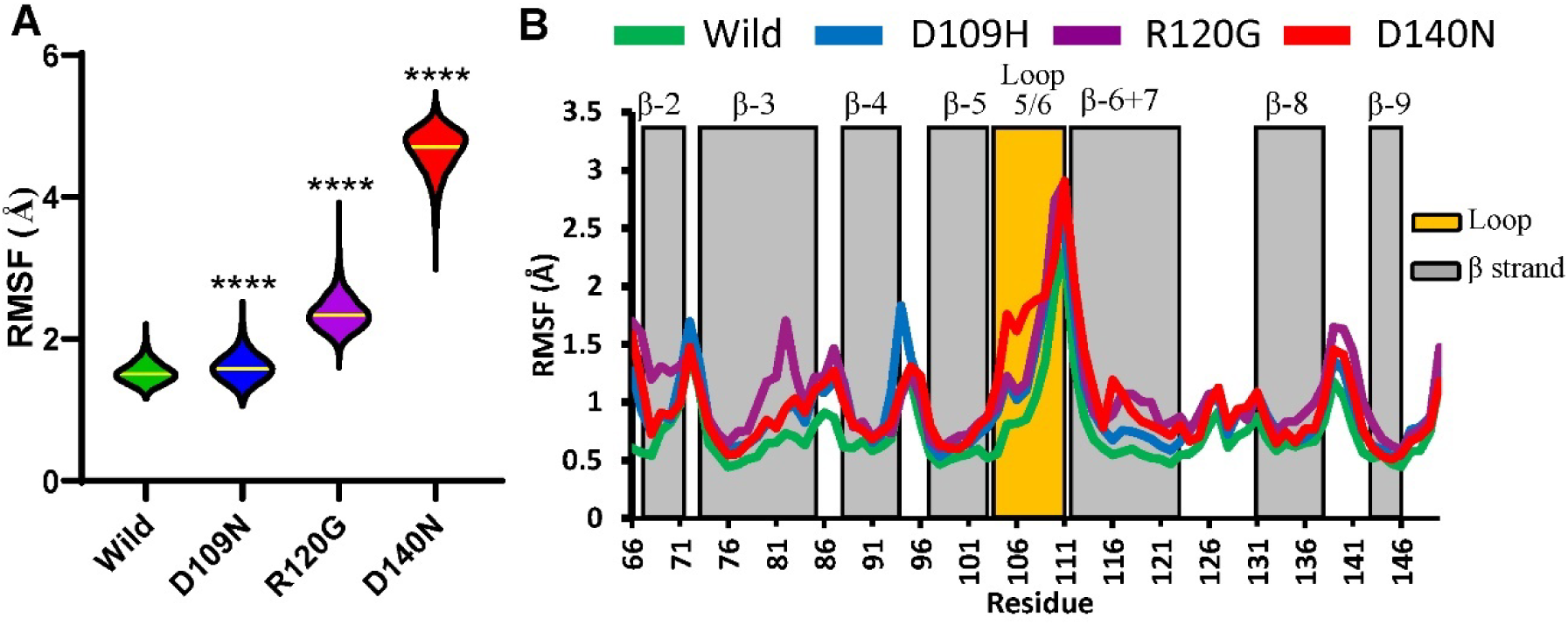
Mutational effects in residual fluctuations. (A) The violin plots indicate each system’s RMSF distribution and the statistical significance considering \*\*\*\**p*<0.05. (B) The root mean square fluctuation (RMSF) graph for both wild and variants. The RMSF of wild, D109H, R120G, and D140N are represented as green, dark blue, purple, and red lines, respectively.

Later on, the dynamic cross-correlation matrix analysis was used to describe the correlative and anti-correlative motions of different systems during the simulation. The DCCM deviation plot is depicted in **Fig. 7A**, where the red colour indicates positive correlations and the blue colour indicates negative correlations. The highly saturated colour represents higher positive or negative correlated motions. The mutant variants showed significantly different positive and negative correlative motions compared to the wild type. The D109H variant showed positive correlative motion between AA_82-90_, AA_98-102_, and AA_137-145_, whereas strong negative correlative motion is shown between AA_107-115_ residues (**Fig. 7Ab**). The R120G variants showed positive correlative motion between AA_90-99_, AA_113-138_, and AA_141-151_ residues, whereas negative correlative motion is shown between AA_90-103_, AA_108-136_, and AA_141-151_ residues (**Fig. 7Ac**). The D140N variants showed more significant and stronger positive and negative correlative motions compared to all variants. It showed strong positive correlative motion between AA_83-91_, AA_99-103_, AA_106-116_, AA_118-130_, AA_135-145_, and AA_147-152_ residues, whereas negative correlative motion is shown between AA_77-95_, AA_97-103_, and AA_125-132_ residues (**Fig. 7Ad**). Moreover, it showed strong negative correlative motion between AA_105-116_, AA_118-124_, AA_135-145_, and AA_147-152_ residues (**Fig. 7Ad**).

**Fig. 7:**
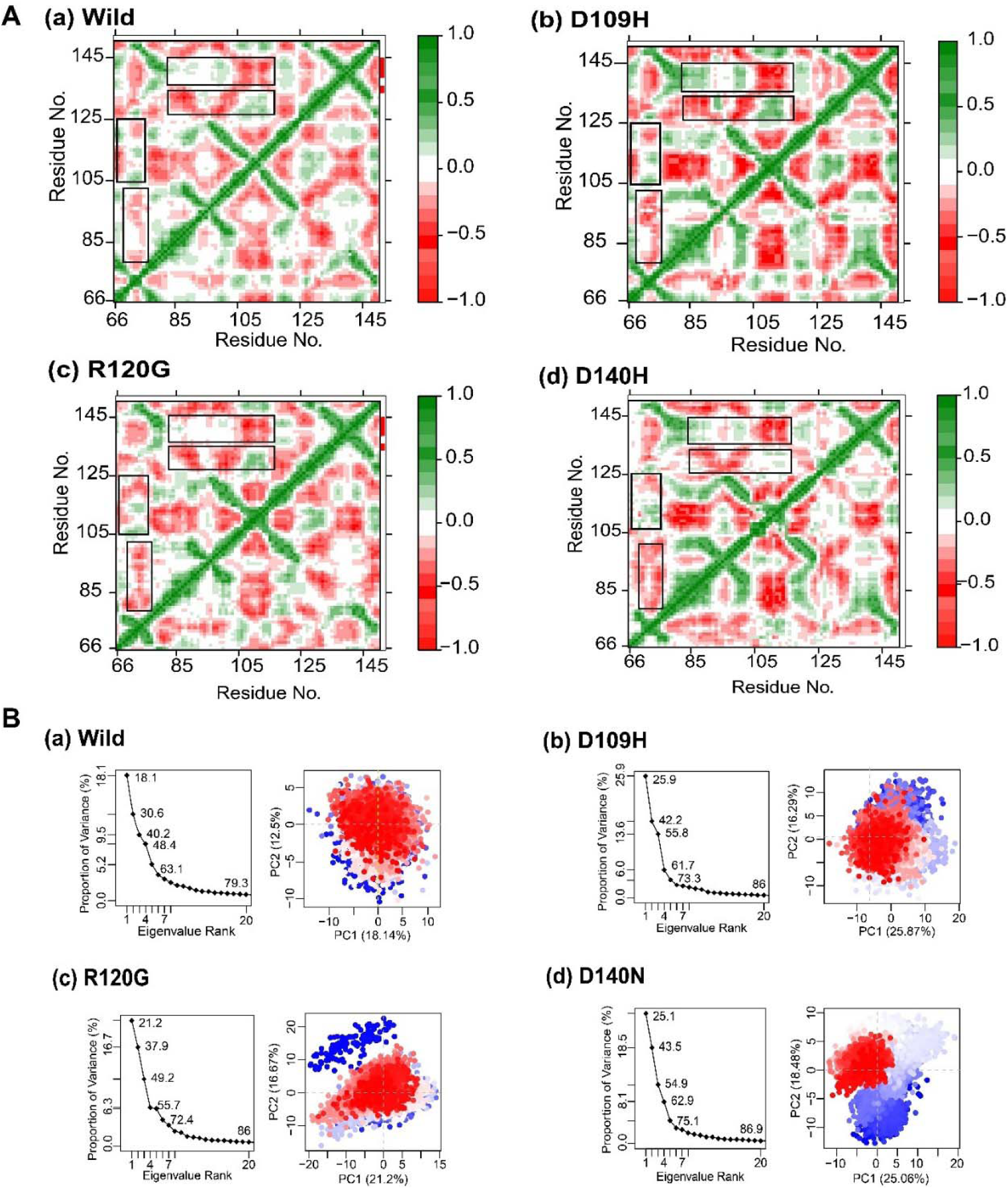
Effects of variants on protein dynamics movement. (**A**) The dynamic cross-correlation maps (DCCMs) of all systems, and (**B**) Principal component analysis of (a) wild, (b) D109H, (c) R120G, (d) D140N in which every single dot defines the protein conformation, where simulation time is characterized by a colour-coded scale from blue to white to red.

In addition, the principal component analysis (PCA) based on Cα atoms was performed to get further insights into protein dynamics. The PCA provides the most significant motions of the residues through the eigenvalues and eigenvectors, whereas the eigenvectors represent atomic motion, and eigenvalues represent atomic contribution in that motion [68]. The calculated PCA values are represented in **Fig. 7B**. The first three PCAs values of wild, D109H, R120G, and D140N, were 18.1, 25.9, 21.2, and 25.1%, respectively (**Fig. 7Bb-d**). The total variance score for wild was 79.3%, while D109H, R120G, and D140N variants provided 86.0, 86.0, and 86.9% scores, respectively (**Fig. 7Ba-d**). This result indicates the increased motion of mutant variants. Fig. **7B** indicated the conformational circulation by blue and red dots, whereas white dots represent the intermediate state. The regular colour schema from red to white and white to blue represents the periodic jumps of conformation. The subspace of PC 1/2 and PC 1/3 of R120G displayed the periodic jumps with significant energy barrier, whereas the wild and D109H and D140N variants were shown a quite similar and overlapping PC subspace without any energy barrier (**Fig. 7B**). The individual residual aberration was also analyzed to differentiate between wild and mutants’ proteins considering PC1 (**Fig. 8**). The major changes occurred at the loop 5/6 and β-6/7 regions. The β-6/7 mediates the dimer interaction of ACD, which is crucial for proper chaperone function. In sum, PCA results support the previous RMSD, Rg, SASA, RMSF, and DCCM results, ensuring the validity of this analysis.

**Fig. 8:**
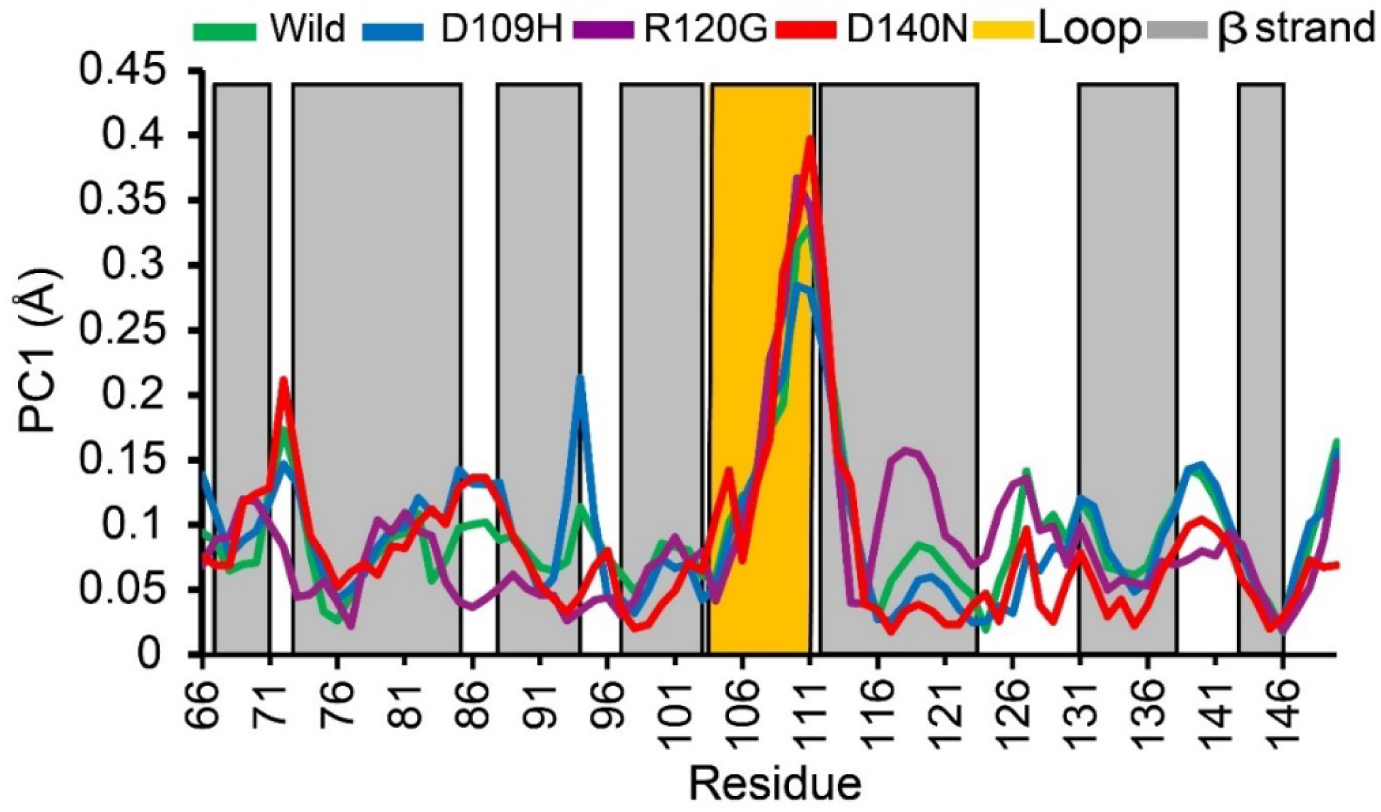
Residue-wise movement plot for PC1, where green, dark blue, purple, and red colour denote wild, D109H, R120G, and D140N, respectively.

## 4 Discussion

The human sHSP HSPB5 is an ATP independent, stress-inducible protein that prevents protein misfolding and aggregation through the interaction with other related proteins [69]. The chaperon activity of HSPB5 improves neurodegenerative disease and other proteotoxicities, including cataracts and different myopathies. As described before, perturbation (mutation, or others) of HSPB5 causes the loss of chaperone activity [19, 22]. It is also evidenced that HSPB5 R120G mutation modulates the dimer interface and hampers the complete oligomeric form of sHSP, which subsequently modulates inter sHSP rearrangement and reduces chaperoning activity [70].

There are various forms of mutations but, among them, nsSNPs have the most significant effect on protein structure and functions [71]. Recently, *in silico* SNP prediction methods gained much attention to reveal the effect of the mutation on protein structures and functions [72–74]. The mutational perturbance of HSPB5 has not been studied yet to elucidate the structural and functional consequences. To deal with this problem, we employed various computational methods, i.e., sequence-based, structure-based, and conservation-based, to emphasize our prediction accuracy [75, 76] along with MD simulation. The advantage of using sequence-based methods is that they can be applied to unknown protein structures, whereas the protein crystal structure is required in structure-based methods [75]. In recent years, the use of multiple *in silico* SNP analysis tools and MD simulation provides the best prediction output, like shown in Anaplastic lymphoma kinase (ALK) [77], Oculocutaneous albinism type III (OCA3) [66], Ataxin-2 (ATXN2) [76], Pyruvate kinase (PK.) M2 [78], Profilin 1 (PFN1) [79], Superoxide dismutase 3 (SOD3) [80], Triggering receptor expressed on myeloid cells 2 (TREM 2) [81], Homogentisate-1, 2-dioxygenase (HGD) [82], and Solute carrier family 26 members 4 (SLC26A4) [83]. However, Brown et al. suggested that at least four or five tools should be considered to predict SNPs’ effect [84]. The statistical correlation analysis depicted that the prediction tools are correlated, rectifying the relevance of considering these tools. We retrieved 313 missense SNPs of HSPB5. This study found that D109H, R120G, and D140N are the most deleterious nsSNPs of HSPB5. The non-synonymous point mutations in conserved regions are more deleterious than the mutation in non-conserved regions [62]. Moreover, the conserved region residues are biologically significant and also crucial for protein-protein interactions [85]. We found that the D109H, R120G, and D140N variants are present in the highly conserved region, specifically in the α c core domain, ACD.

MD simulation provides detailed atomic insight, including the structural consequences of any perturbation with a finite resolution. Besides, MD simulation allows the study of the conformational transformation and offers relations between protein structure and dynamics [86]. The dynamic nature of protein is programmed in its structure and mostly vital for functions [86]. The local structural fluctuations may lead to higher structural deviation in RMSD, even though their global topologies are very close [80]. The higher RMSD fluctuations represent lower stability [81, 87]. The mutant variants showed higher RMSD values than the wild, which depicts their higher flexibility (**Table 2 and Fig. 4**). Rg is generally defined as the average mass-weighted root distance for a particular group of atoms from their common center of mass so that the Rg values are directly related to the volume of the protein. This analysis, therefore, represents the detailed structural dimensions [88]. The higher Rg values of D109H, R120G, and D140N indicate that mutation reduced the compactness. The detailed protein interaction profile is calculated using SASA. SASA is directly proportional to its interaction ability [88]. It indicates that the studied variants D109H and R120G are comparatively more accessible than the wild type protein, and D140N is less accessible, which possibly will modulate the capacity of these variants to interact with other oligomers [80]. The conformational drifting represented in the Rg (**Fig. 5A**) and SASA (**Fig. 5B**) calculation indicated that the studied mutation could affect the protein interaction capability [89]. We found that wild protein has fewer Rg values than the mutants (**Table 2**), which evidenced that D109H, R120G, and D140N reduce the stability of protein and hereafter decrease the functionality of the protein [80]. The flexibility of protein is directly proportional to the number of intermolecular H-bonds between the amino acid residues [90, 91]. The reduction of the number of H-bonds discussed previously was validated by estimating the protein’s intramolecular hydrogen bond’s interaction (**Table 2**). To get the insights of flexibility in mutant D109H, R120G, and D140N variants, we plotted a graph considering the change in the number of H-bonds over time (**Fig. 5C**). The average number of H-bonds was less in mutants compared to the wild protein. The mutant structure shows fewer H-bonds due to their flexibility (**Fig. 5C**). This result suggests that wild residue substitution might increase the RMSF fluctuations, as shown in **Fig. 6** [92]. Moreover, H-bonds define protein structure, folding, and recognition as essential for their functionality [93]. The mutant variants’ average H-bonds were decreased compared to the wild protein (**Table 2**), representing the conformational change of the variants.

The fluctuated RMSD values make sense to analyze the local residual fluctuations; thus, we calculated RMSF [94]. As protein flexibility seems to have a significant effect on the binding thermodynamics and can support or deter interaction [95, 96], the flexibility of variations is shown during RMSF (**Fig. 6**) calculation represented that the D109H, R120G, and D140N mutations could distress the αB-c ability to interact with other sHSP. According to the RMSD, Rg, SASA, RMSF analyses, the mutant variants showed higher flexibility. We found that the D140N variant is more harmful in terms of protein flexibility, which might prime distress the functional behaviour of protein and reduce chaperoning activity. In a natural setting, proteins continuously change their conformation for a particular function, and their residual correlative and anti-correlative motions play a significant role in substrate management. However, mutational perturbance hampers this communication [97]. The mutant variants showed significant positive and negative correlative motions compared to the wild protein, indicating their conformational shifting that may cause loss of function. The higher anti-correlative motions of these mutant variants indicate their conformational flexibility and functional disturbance (**Fig. 7A**), which might be noteworthy to cut chaperoning activity. Furthermore, the systems’ collective motion dynamics were analyzed using PCA. PCA describes the systems’ overall mutational effects and mechanical features [41]. PCA analysis represents the leading modes in the motions of the molecules [98, 99]. Thus, it is often used to understand the conformational drifting in protein folding [100] and structural dynamics influenced by mutation [41, 101]. Strongly, the analyzes of PCA and RMSF produce the same structural dispersion attributes of systems.

The mutant’s variants’ conformations were stretchy during the simulation period resulting in a prolonged unstable conformation than the wild protein. The change in amino acid disturbs the protein domain stability and function, which leads to the modification of the mutants’ structural dynamics. The protein’s conformational stability is solely related to the pathogenic mutations of individuals. It is therefore shown that mutations affect the protein compactness that eventually leads to destabilization and dysfunction. Besides, exposure to protein hydrophobic residues could also provoke misfolding, and here change of arginine (R120) to glycine might have a higher chance to basket native folding pattern. Moreover, this conformational change could drive the “guilt-by-association” mechanism to lose the PN network functions. Therefore, the mentioned genomic change could circuitously compromise the arrangement and role of various cellular proteins associated with proteostasis.

## 5 Conclusion

The present study has predicted damaging SNPs in the *CRYAB* (HSPB5) gene with the assistance of several cutting-edge *in silico* tools and analyzed their structural and functional effects on HSPB5 (αB-c) protein. The findings clearly showed the three most dangerous SNPs, i.e., D109H, R120G, and D140N, at highly conserved ACD domain of HSPB5 while dynamics simulation studies represent the mutational destabilization of ACD. Compared to the wild type, the R120G and D140N mutations showed more structural variations associated with the increased protein flexibilities and significant structural drifting. The structural shifting might alter the chaperoning activity of HSPB5. This study offers a possible arena to design therapeutics to treat patients with proteostasis imbalance resulting from these mutations. However, further research is required to investigate more insights into these variants and their implications in associated diseases.

## Author Contributions

MSK, MCA contributed to research designing, manuscript writing, data curation & analysis, table, and figure processing. MCA and RD conducted the whole experiments. MSK, LB and MSK contributed to the initial SNP analysis. YAM contributed to initial simulation data processing. RD and MMR contributed to research planning, supervision, and critical revision. All authors read and approved the final version of the manuscript.

## Acknowledgment

We are thankful to Professor Dr. Il Soo Moon, Department of Anatomy, Dongguk University College of Medicine, Gyeongju 38066, Republic of Korea, for providing high-end computer access to complete this project. We are also grateful to Zulkar Nain, Department of Biotechnology and Genetic Engineering, Faculty of Biological Sciences, Islamic University, Kushtia 7003, Bangladesh, and Md Rimon Parves, Department of Biochemistry and Biotechnology, University of Science and Technology Chittagong (USTC), Foy’s Lake, Khulshi, 4202, Chittagong, Bangladesh for their enormous suggestions to improve the quality of this manuscript.

